# Liver Electrical Impedance Tomography for Early Identification of Fatty Infiltrate in Obesity

**DOI:** 10.1101/2020.12.21.423854

**Authors:** Chih-Chiang Chang, Zi-Yu Huang, Shu-Fu Shih, Yuan Luo, Arthur Ko, Qingyu Cui, Susana Cavallero, Swarna Das, Gail Thames, Alex Bui, Jonathan P. Jacobs, Päivi Pajukanta, Holden Wu, Yu-Chong Tai, Zhaoping Li, Tzung K. Hsiai

## Abstract

Non-alcoholic fatty liver disease (NAFLD) is endemic in developed countries and is one of the most common causes of cardiometabolic diseases in overweight/obese individuals. While liver biopsy or magnetic resonance imaging (MRI) is the current gold standard to diagnose NAFLD, the former is prone to bleeding and the latter is costly. We hereby demonstrated liver electrical impedance tomography (EIT) as a non-invasive and portable detection method for fatty infiltrate. We enrolled 19 subjects (15 females and 4 males; 27 to 74 years old) to undergo liver MRI scans, followed by EIT measurements via a multi-electrode array. The liver MRI scans provided subject-specific *a priori* knowledge of the liver boundary conditions for segmentation and EIT reconstruction, and the 3-D multi-echo MRI data quantified liver proton-density fat fraction (PDFF%) as a recognized reference standard for validating liver fat infiltrate. Using acquired voltage data and the reconstruction algorithm for the EIT imaging, we computed the absolute conductivity distribution of abdomen in 2-D. Correlation analyses were performed to compare the individual EIT conductivity vs. MRI PDFF with their demographics in terms of gender, BMI (kg·m^−2^), age (years), waist circumference (cm), height (cm), and weight (kg). Our results indicate that EIT conductivity (S·m^−1^) and liver MRI for PDFF were not correlated with the demographics, whereas the decrease in EIT conductivity was correlated with the increase in MRI PDFF (*R* = − 0.69, *p*= 0.003). Thus, EIT conductivity holds promise for developing a non-invasive, portable, and quantitative method to detect fatty liver disease.

## Introduction

Obesity is the major risk factor associated with the development of nonalcoholic fatty liver disease (NAFLD), affecting more than a third of American. adults, and the prevalence of severe obesity (BMI ≥ 35 kg·m^−2^) is continuing to rise nationwide^1^. NAFLD is now one of the most common causes of cirrhosis requiring liver transplantation in the Western world^2,3^. A clinical challenge in the management of NAFLD resides in non-invasively detecting fatty liver (i.e., simple hepatic steatosis) and monitoring disease progression to steatohepatitis (hepatic inflammation), fibrosis (liver scarring), and ultimately cirrhosis^4,5^. While liver biopsy remains the gold standard for diagnosis of NAFLD, it carries a substantial risk of bleeding and is confounded by sampling bias and inter-observer variability^6^. While liver MRI proton-density fat fraction (PDFF) is recognized as the non-invasive reference standard for validating liver fat infiltrate^7,8^, it is costly for underserved communities. While ultrasound is non-invasive, it is limited by spatial resolution and operator dependency ^9,10^. Thus, there remains an unmet clinical need to develop a non-invasive and economic method that is operator-independent and portable for early detection of fatty liver disease.

To this end, we demonstrated in prior study the theoretical and experimental basis of electrical impedance tomography (EIT) for measuring liver fat content in the New Zealand White Rabbit model of atherosclerosis and fatty liver disease^11^. By virtue of tissue-specific electrical conductivity, fatty infiltrate in the liver was characterized by its frequency-dependent electrical impedance (Z) in response to applied alternating current (AC)^11^. At low frequencies, the cell membranes impede the current flow, resulting in high conductivity, whereas at high frequency, they serve as the imperfect capacitors, resulting in tissue- and fluid-dependent impedance. This impedimetric property enables the development of a multi-electrode array to measure tissue-specific conductivity, morphology, and changes in 3-D volume in response to changes in cardiac output or lung capacity^12–15^. A host of literature has demonstrated the application of EIT for functional studies of the brain, cardiac stroke volume, and respiratory ventilation (transthoracic impedance pneumography)^16–19^.

In this context, we reconstructed liver frequency-dependent conductivity distribution with the multi-electrode array-acquired voltage data to demonstrate liver EIT.^11^ Applying the multi-electrode array, we performed EIT voltage measurements by biasing electrical currents (at 2-4 mA and 50-250 kHz) to the upper abdomen. The currents penetrated the body to varying depths, and the resulting boundary voltages were acquired by the electrodes. In response to the applied alternating current (AC), muscle and blood are more conductive than fat, bone, or lung tissue due to the varying free ion content.^20,21^ Fat-free tissue such as skeletal muscle carries high water (~73%) and electrolyte (ions and proteins) content, allowing for efficient electrical conductivity (S·m^−1^), whereas fat-infiltrated tissue such as fatty liver (steatosis)^22^ is anhydrous, resulting in a reduction in conductivity^23^. This impedimetric property provides the theoretical principle to apply the portable liver EIT for the early identification of fatty liver infiltrate with translational implications for the prevention of liver fibrosis and major adverse coronary events (MACE). Unlike EIT for cardiopulmonary function focusing on the differential conductivity ^12–15, 16–19^, we addressed the ill-posed inverse problem to demonstrate the absolute liver conductivity in 2-D.

We recruited overweight/obese subjects to undergo liver 3T MRI scans, followed by voltage measurements via the flexible multi-electrode array (Swisstom AG, Switzerland) for EIT. MRI images were acquired to provide subject-specific *a priori* knowledge of the liver geometry for performing liver segmentation and positioning to solve the inverse problem of EIT reconstruction. We further compared the subject-specific EIT conductivity with the liver MRI proton-density fat fraction PDFF as a reference standard for validating fatty liver infiltrate^24^. Next, we performed the Pearson’s correlation analyses between the EIT liver conductivity and demographic parameters, and also performed the correlation analyses between MRI PDFF and demographics parameters in terms of gender, BMI (kg·m^−2^), age (years), waist circumference (cm), height (cm), and weight (kg). Following Bonferroni correction for multi-testing, correlation analyses revealed that liver EIT conductivity (S·m^−1^) and MRI PDFF were not correlated with these demographics; however, the EIT liver conductivity map was negatively correlated with the MRI PDFF. This inverse correlation between the EIT liver conductivity and MRI PDFF holds promises for developing non-invasive and portable liver EIT for early detection of silent fatty liver content in the healthy overweight/obese individuals

## Results

### Schematic workflow illustrates the steps to compare and validate the EIT reconstruction with MRI

The recruitment of subjects followed the guidelines of the Human Subjects Protection Committee of UCLA, described in the method section. For the workflow and schematic setup (**Fig 1**), each subject would undergo a different series of MRI scans, including a 30-min MRI multi-echo imaging to acquire the PDFF map of the liver. Next, EIT measurement with 32 electrodes attached to the subject’s abdominal region was performed right after the MRI scan to obtain the corresponding liver fat measurement. The average PDFF of the liver, as well as the EIT conductivity of the liver, were then quantified for validation and comparison.

**Figure 1.**
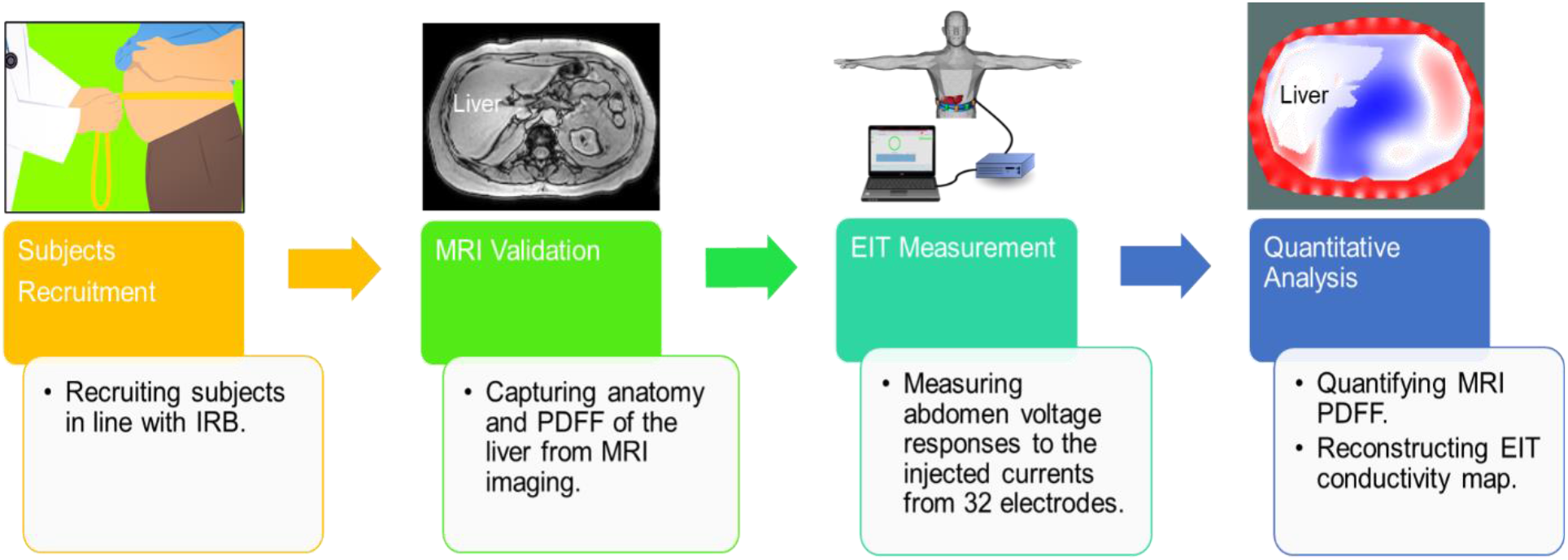
Schematic workflow of the comparison and validation of the MRI and EIT. Volunteers were recruited in line with the UCLA Institutional Human Subjects Protection Committee. Multi-echo MRI scans were performed to provide the liver anatomy and proton density fat fraction (PDFF), followed by the EIT measurements. Finally, the EIT conductivity maps were reconstructed and the MRI PDFF was used to quantify fatty liver infiltrate and to compare with EIT liver conductivity.

### Comparison between MRI multi-echo imaging and EIT images

Liver MRI images provide *a priori* geometric information to reconstruct 2D EIT images. This information includes the boundary conditions for: 1) the abdominal cross-section, 2) the peripheral tissues consisting of the skin, subcutaneous fat, and the ribs, and 3) the liver in the abdomen. With this information, liver EIT inverse problem was solved as described in the Methods section. For each subject, EIT liver conductivity and MRI PDFF were compared with the corresponding BMI value (**Table 1**). Also, the representative abdomen MRI images for anatomy and PDFF, liver segmentation (annotation), and liver EIT conductivity maps (S·m^−1^) were compared (**Fig. 2**). We observed that the MRI PDFF and EIT liver conductivity were not correlated with the magnitude of BMI. Despite a negative correlation with EIT, the MRI PDFF for Subject 17 with a relatively lower BMI (BMI = 27.1 Kg·m^−2^, PDFF = 6.2%, EIT = 0.3243 S·M^−1^) was higher than that of Subject 3 with a high BMI value (BMI = 39.0 Kg·m^−2^, PDFF = 3.82%, conductivity = 0.3296 S·M^−1^). We also noted that MRI PDFF for Subject 11 with a low BMI value (BMI = 27.9 Kg·m^−2^, MRI PDFF = 3.62%, EIT = 0.3473 S·M^−1^) was lower than that of Subject 10 with a high BMI values (BMI = 34.3 Kg·m^−2^, MRI PDFF = 16.44%, EIT = 0.3007 S·M^−1^). Despite similar BMI (27.1 Kg·m^−2^ vs. 27.9 Kg·m^−2^), the percentages of MRI PDFF of Subject 17 was around two times higher than that of Subject 11 (6.20 vs. 3.62 %). Notably, the MRI PDFF for Subject 10 (BMI = 34.3 Kg·m^−2^) was more than 4 times higher than that of Subject 3 (BMI = 39.0 Kg·m^−2^). These inconsistent comparisons suggest that BMI may not be the ideal index for predicting the levels of fatty liver infiltrate, and Pearson’s correlation analyses were performed in the ensuing results.

**Table 1.** List of BMI (Kg·m^−2^), MRI PDFF (%) and EIT liver conductivity of all subjects. (S·M^−1^).

| Subjects | BMI(Kg.m<sup>-2</sup>) | EIT (S.m<sup>-1</sup>) | MRI PDFF (%) |
| --- | --- | --- | --- |
| 1 | 34.4 | 0.3518 | 2.14 |
| 2 | 49.7 | 0.3290 | 4.05 |
| 3 | 39.0 | 0.3296 | 3.82 |
| 4 | 33.0 | 0.3819 | 27.89 |
| 5 | 30.6 | 0.3377 | 10.51 |
| 6 | 36.3 | 0.3444 | 4.14 |
| 7 | 29.3 | 0.3280 | 2.41 |
| 8 | 37.8 | 0.3405 | 2.25 |
| 9 | 32.0 | 0.3381 | 6.53 |
| 10 | 34.3 | 0.3007 | 16.44 |
| 11 | 27.9 | 0.3473 | 3.62 |
| 12 | 46.8 | 0.3307 | 5.14 |
| 13 | 38.9 | 0.3305 | 3.31 |
| 14 | 25.5 | 0.3010 | 2.11 |
| 15 | 33.7 | 0.3306 | 10.78 |
| 16 | 27.4 | 0.3407 | 1.08 |
| 17 | 27.1 | 0.3243 | 6.20 |
| 18 | 46.9 | 0.3455 | 18.56 |
| 19 | 29.8 | 0.3507 | 2.29 |

**Figure 2.**
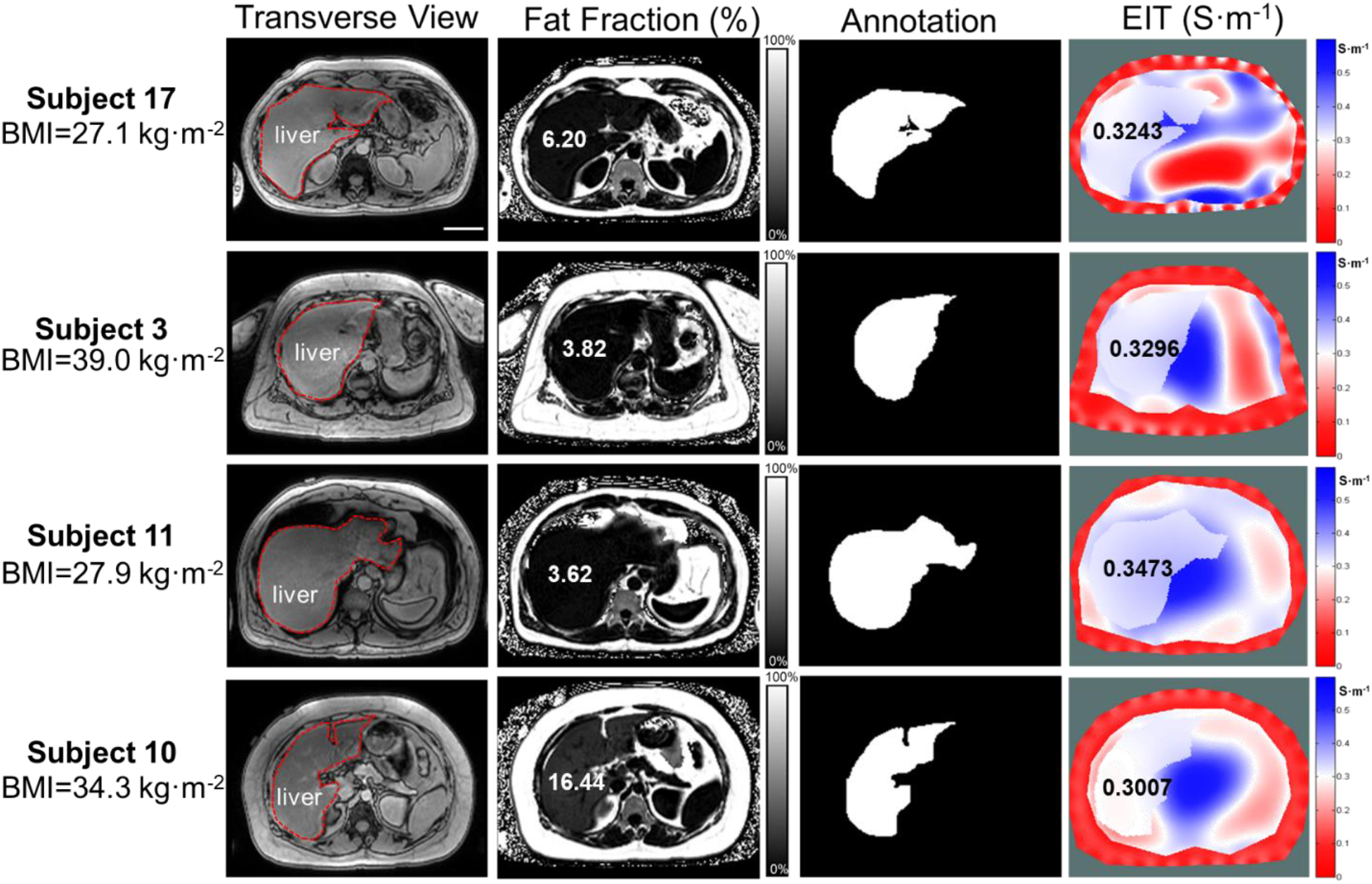
Representative MRI multi-echo and EIT images. Four representative subjects with different BMI values (Kg·m^−2^) were compared with MRI PDFF (%) and EIT conductivity (S·M^−1^), respectively. The transverse MRI views demarcate the liver anatomy, the fat fractions provide the corresponding MRI PDFF, annotation reveals the liver boundary condition following image segmentation, and 2-D EIT images unveil the abdomen conductivity distribution and average liver conductivity. The subject 17 with a BMI of 27.1 Kg·m^−2^ developed a relatively high MRI PDFF (6.2%) and a low EIT liver conductivity (0.3243 S·M^−1^); whereas the subject 3 with BMI of 39 Kg·m^−2^ developed a relatively low MRI PDFF (3.82%) and high EIT liver conductivity (0.3296 S·M^−1^). However, the subject 11 with BMI of 27.9 Kg·m^−2^ developed a relatively low MRI PDFF (3.62%) in association with a relatively high EIT liver conductivity (0.3473 S·M^−1^), and the subject 10 with BMI of 34.3 Kg·m^−2^ also developed a relatively high MRI PDFF (16.44%) in association with a low EIT liver conductivity (0.3007 S·M^−1^). These initial comparisons suggest inconsistent correlations between BMI and MRI PDFF and EIT liver conductivity. Scale bar: 8 cm.

### EIT conductivity vs. MRI PDFF

Using the MRI PDFF and EIT conductivity data from **Table 1**, we performed the Pearson’s correlation analyses and identified whether the magnitude of BMI correlates with the percentage of MRI PDFF or EIT conductivity. We demonstrated that the correlation between BMI and MRI PDFF (*R* = −0.037, *p*=0.89, n= 16) and the correlation between BMI and EIT (*R* = −0.19, *p* = 0.47, n= 16) were statistically insignificant (**Fig. 3A-B**). However, the confidence interval plot revealed statistically significant negative correlation between EIT and MRI PDFF (*R* = −0.69, *p* = 0.003, n= 16) (**Fig. 3C**). This finding suggests that EIT conductivity may be used as an index for non-invasive detection method to quantify human liver fatty infiltrate.

**Figure 3.**
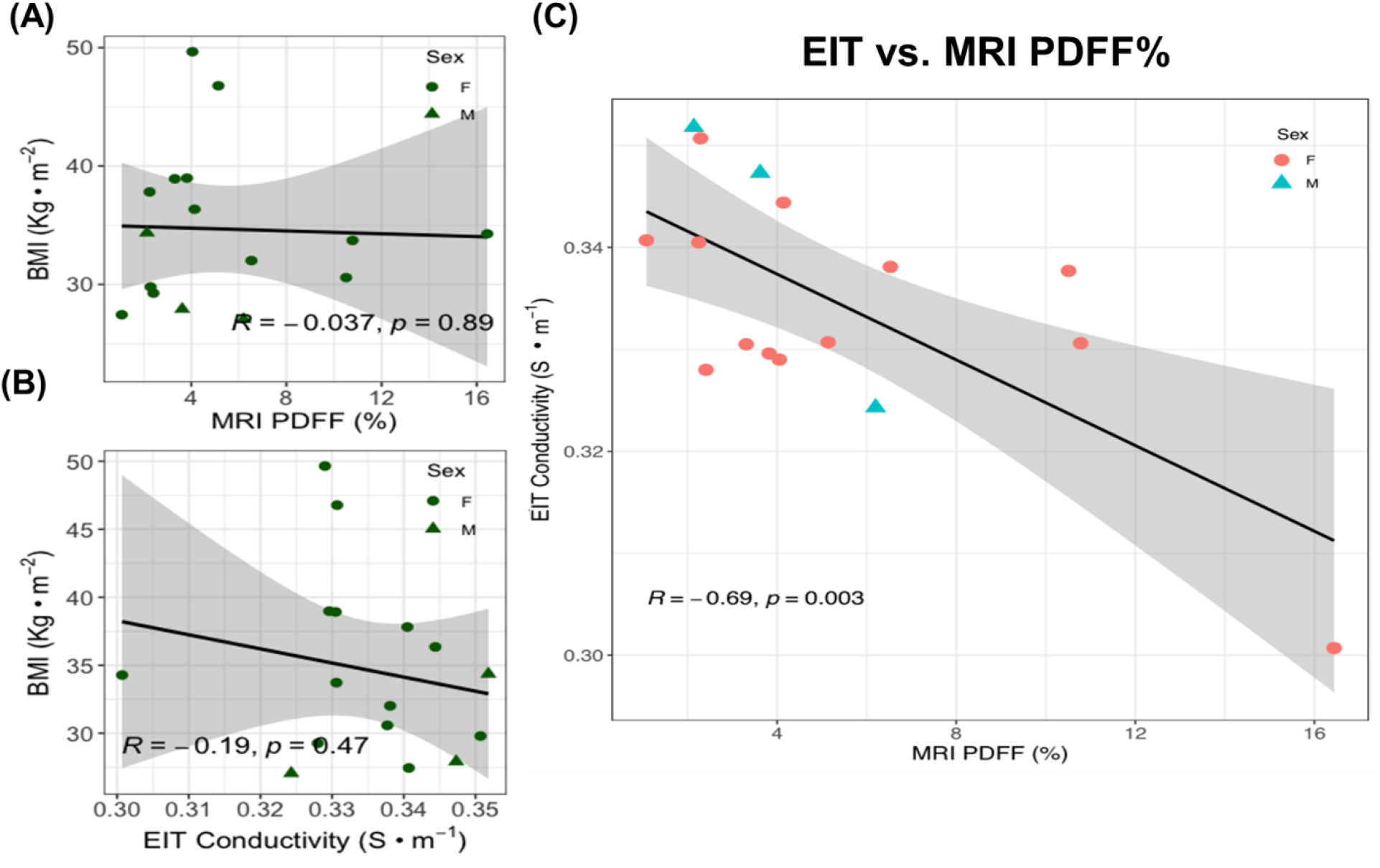
Statistical analyses of BMI vs. MRI PDFF and vs. EIT liver conductivity. **(A)** BMI values are not correlated with MRI PDFF. (Pearson correlation coefficient *R* = − 0.037, *p* = 0.89, n=16). **(B)**. BMI values were also not correlated with EIT liver conductivity values (*R* = − 0.037, *p* = 0.89, n=16). **(C)** EIT liver conductivity values were negatively correlated with MRI PDFF (R = − 0.69, *p* = 0,003, n=16). The shaded areas reflect the 95% confidence intervals of the linear slopes.

### Correlation analyses with the demographic parameters, MRI PDFF, and EIT conductivity

To demonstrate liver EIT for identification of fatty liver infiltrate in the enrolled subjects (BMI > 25), we performed correlation analyses with demographic parameters including age, waist circumference, height, and weight, respectively (**Table 2**). We compared the correlation coefficients between MRI PDFF and the demographic parameters (**Fig. 4**). Following the Bonferroni correction for multi-testing, the correlations with age (*R* = −0.13, *p* = 0.64, n= 16), waist circumference (*R* = −0.23, *p* = 0.4, n= 16), height (*R* = −0.59, *p* = 0.016, n= 16) and weight (*R* = −0.41, *p* = 0.12, n= 16) were statistically insignificant. We further compared the correlation coefficients between liver EIT and demographic parameters (**Fig. 5**). The correlation with age (*R* = −0.1, *p* = 0.71, n= 16), waist circumference (*R* = −0.05, *p* = 0.85, n= 16), height (*R* = −0.63, *p* = 0.0092, n= 16) and weight (*R* = −0.19, *p* = 0.47, n= 16) were statistically insignificant. Thus, these analyses corroborate that BMI and other demographic parameters were not correlated with liver fat infiltrate in our overweight/obese subjects.

**Table 2.** Demographics of overweight/obese subjects. The demographics of 19 subjects including sex, BMI, age, waist circumference, height, and weight, are demonstrated.

| Subjects | Sex | BMI(Kg·m<sup>-2</sup>) | Age (year) | Waist Circumference (cm) | Height (cm) | Weight (kg) |
| --- | --- | --- | --- | --- | --- | --- |
| 1 | M | 34.4 | 41 | 116 | 175 | 105.2 |
| 2 | F | 49.7 | 67 | 131 | 158.5 | 124.7 |
| 3 | F | 39.0 | 63 | 123.5 | 160 | 99.8 |
| 4 | F | 33.0 | 35 | 116.5 | 168.9 | 94.1 |
| 5 | F | 30.6 | 61 | 92.5 | 155.5 | 73.9 |
| 6 | F | 36.3 | 27 | 103.5 | 163.5 | 97.2 |
| 7 | F | 29.3 | 42 | 91 | 155 | 70.3 |
| 8 | F | 37.8 | 60 | 115.5 | 174 | 114.5 |
| 9 | F | 32.0 | 36 | 113 | 170 | 92.5 |
| 10 | F | 34.3 | 36 | 101.5 | 152 | 79.2 |
| 11 | M | 27.9 | 47 | 95 | 178 | 88.5 |
| 12 | F | 46.8 | 39 | 130 | 160 | 119.8 |
| 13 | F | 38.9 | 48 | 114 | 168 | 109.9 |
| 14 | F | 25.5 | 74 | 96 | 163.5 | 68.2 |
| 15 | F | 33.7 | 26 | 95 | 153 | 78.9 |
| 16 | F | 27.4 | 33 | 93.5 | 170.5 | 79.8 |
| 17 | M | 27.1 | 47 | 103 | 178 | 85.7 |
| 18 | M | 46.9 | 57 | 141.5 | 177.5 | 147.7 |
| 19 | F | 29.8 | 30 | 102 | 180.3 | 96.9 |

**Figure 4.**
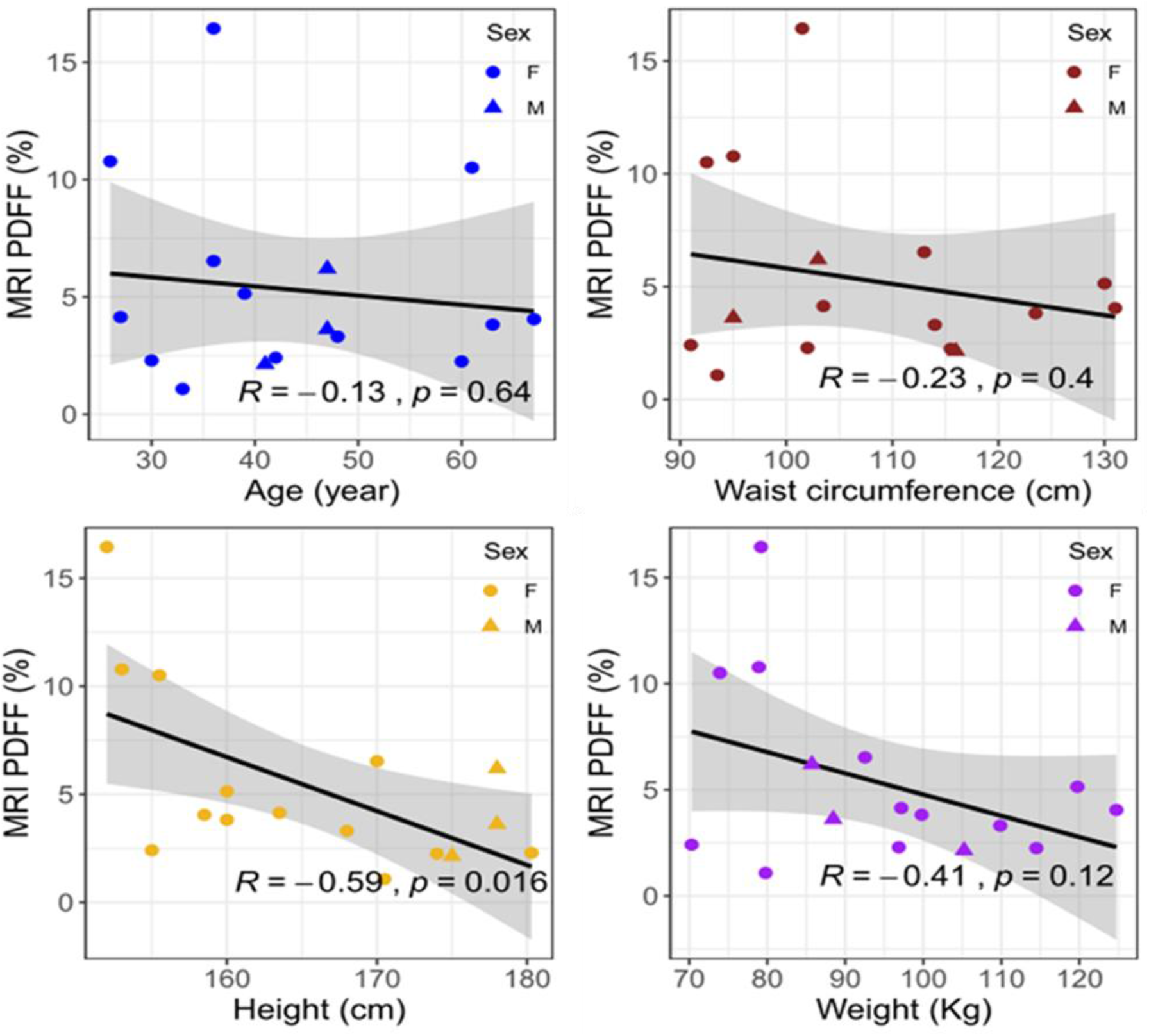
MRI PDFF vs. age, waist, height, and weight. The Pearson correlation coefficients (*R*) and *p* values were analyzed for age, waist circumference, height, and weight, respectively. The circles denote female subjects and triangles denote male subjects. The 95% confidence intervals of the linear slopes are illustrated as shaded area. *R* values are −0.13 for age (*p* = 0.64, n= 16), − 0.23 for waist circumference (*p*= 0.4, n=16), 0.59 for height (*p*= 0.016, n= 16)., and −0.41 for weight (*p*= 0.12, n= 16), demonstrating low to intermediate correlation with MRI PDFF.

**Figure 5.**
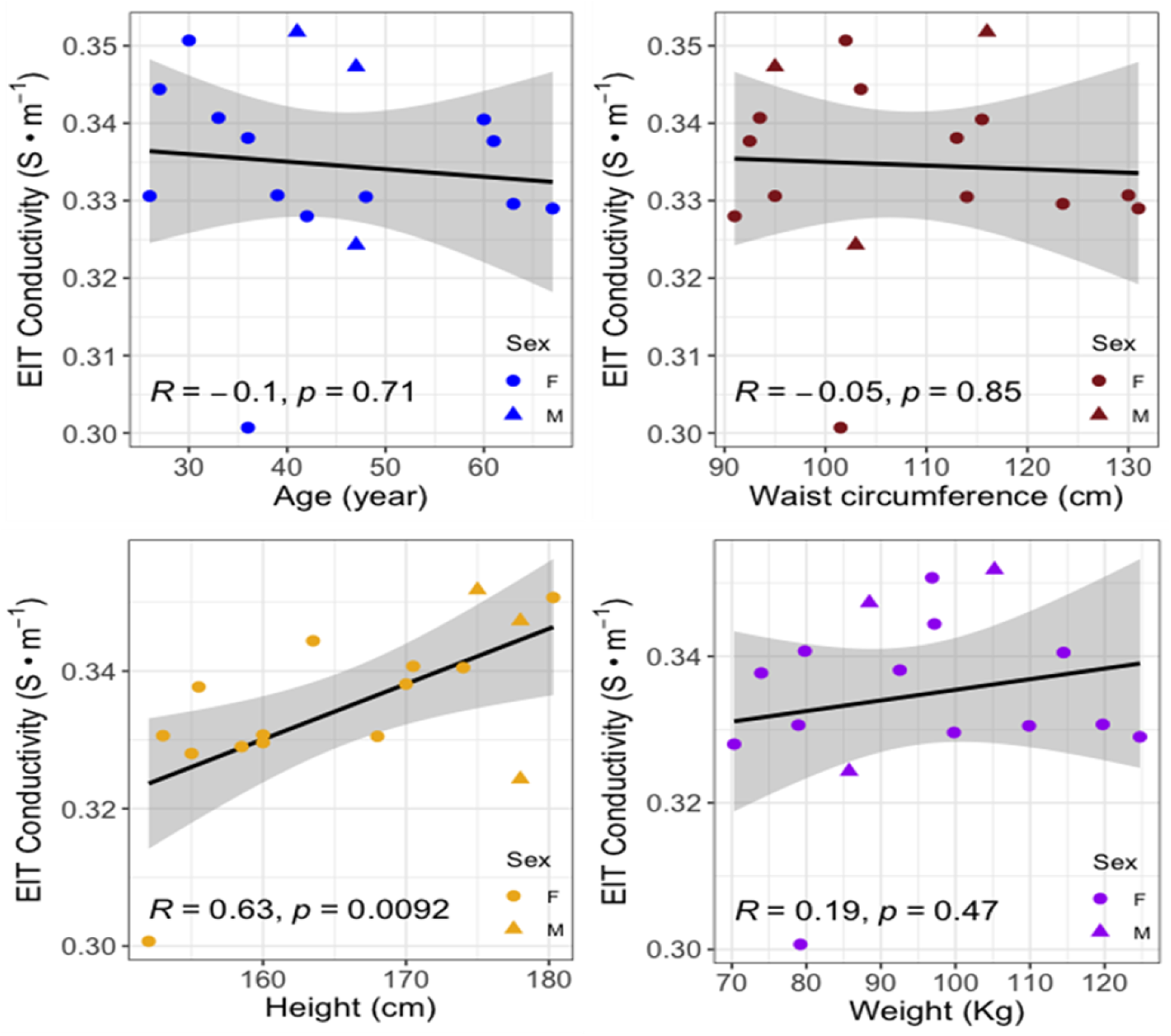
EIT liver conductivity vs. age, waist, height, and weight. The *R* values for age (R = − 0.1, *p* = 0.71, n= 16), waist circumference (R = − 0.05, *p*= 0.85, n=16), height (R = − 0.63, *p* = 0.0092, n=16) and weight (R = −0.19, *p* = 0.47, n=16) demonstrate low to intermediate correlation with EIT conductivity.

## Discussion

Non-invasive and cost-effective monitoring of fatty liver disease remains an unmet clinical need for the early identification of cardiometabolic disorders. While liver biopsy or magnetic resonance imaging (MRI) have been performed to detect non-alcoholic fatty liver disease (NAFLD), the risk of complications, sampling errors and cost limit its clinical application for the general population. We hereby demonstrated the development of liver EIT as a non-invasive and portable detection method for quantifying liver fat content. We recruited 19 overweight/obese adults with BMI > 25 Kg·M^−2^ to undergo liver MRI scans. We performed the individual liver EIT measurements with a multi-electrode array, and we used the anatomic information to address the inverse problem for reconstructing the subject-specific EIT conductivity map. We performed correlation analyses on liver EIT vs. MRI PDFF in relation to the individual demographics^25^. To our best knowledge, this is the first demonstration of statistically significant negative correlation between EIT-acquired liver conductivity and MRI-quantified PDFF.

EIT has been applied to clinical medicine over the past two decades. Diagnostic EIT was developed for pulmonary function and lung capacity^23^. For instance, respiratory monitoring was exhibited by transthoracic impedance pneumography^26,27^, and the cardiac output (CO) and stroke volume (SV) measurements were demonstrated via myocardial motion and blood volume,respectively^28,29^. EIT has also been applied for assessing conductivity in breast tissue and brain^30^. Using the multi-electrode configuration, we obtained voltage from the surface of the abdomen by injection of AC current, to reconstruct the EIT conductivity map inside the liver. While EIT has been extensively studied,^31–37^ the nonlinear forward and inverse models for reconstructing the EIT conductivity map remain a computational challenge. The ill-posed inverse problem introduces issues of existence, uniqueness, and instability of the solution^36^. The non-linear inverse model for EIT reconstruction requires *a priori* knowledge of the anatomic boundaries to enhance the spatial resolution for establishing the absolute conductivity value^38^. To improve the EIT reconstruction, investigators have integrated EIT with other imaging modalities, including co-registration with MRI^39–41^ and introduction of ultrasonic vibration to the target tissue in the presence of the magnetic field. This integration could generate inductive currents within the liver resulting in higher spatial resolution, thus obviating the need for *a priori* knowledge of the object geometry and location needed for EIT reconstruction^42^.

Researchers have also applied other approaches to enhance the algorithm for solving the ill-posed inverse problem. For instance, particle swarm optimization (PSO) was applied via paradigm shift from the conventional Gauss-Newton methods for fast convergence and high spatial resolution to solve EIT^43,44^. Recent studies have applied machine learning, including Convolutional Neural Networks, to solve the non-linear ill-posed inverse problem for accurate EIT reconstruction^45,46^. Hamilton *et al.* obtained absolute EIT images by combining the D-bar method with subsequent processing using convolutional neural networks (CNN) technique for sharpening EIT reconstruction^45^. Li *et al.* utilized deep neural networks (DNN) to directly obtain a nonlinear relationship between the one-dimensional boundary voltage and the internal conductivity^46^. Experimentally, the accuracy of EIT reconstruction may be further enhanced by increasing the electrode arrays at multiple levels around the abdomen. This multi-level configuration would be able to inject the currents to and record the voltages from the entire liver. As a result, 3-D reconstruction of a liver conductivity distribution would be realized.

As a corollary, we compared the liver anatomy with MRI PDFF distributions from a representative 3-D rendering (**Fig S1A-B**). The 3-D EIT conductivity map was reconstructed with the aid of the MRI multi-echo sequence as *a priori* knowledge (**Fig S1C**). The high-fat region in the MRI PDFF map (red dashed box) was also detected by the EIT demonstrating lower conductivity. The 3-D MRI and EIT analyses further support the negative correlation between MRI PDFF and EIT liver conductivity. The 3-D EIT conductivity map reveals the inhomogeneous fat distribution as supported by the MRI 3-D rendering images (**Fig S1C**). With additional scanning along the z-direction, a precise mapping could be reconstructed to unveil the details of the heterogeneous fat distribution.

While MRI images provided the *a priori* knowledge to solve the ill-posed inverse problem for EIT reconstruction, alternative methods to provide such information would allow for low-cost liver EIT screening for the general population. A strain-displacement conversion method for reconstructing the deformed shape of the object with the boundary conditions was proposed by Luo *et al* ^47^. This method would potentially provide the outer abdomen boundary information by embedding the positional sensors in the EIT sensor belt. Another method is to apply differential EIT, which has the potential to identify the tissue-specific conductivity. If the two frequencies are correctly selected, it is possible to differentiate the fatty tissue from the non-fatty tissues or organs by virtue of fatty tissue-specific electrical properties are distinguished from other tissues or organs (**Table S1**) ^48,49^ thus providing the peripheral layer boundaries information. Furthermore, using a large number of MRI image database, we may correlate the liver and peripheral layer boundaries with the waist circumferences. This correlation would provide a calibration curve between the boundary conditions and a demographic parameter. Thus, the aforementioned methods would be the future area of research to obviate the need for the MRI-acquired *a priori* knowledge to improve EIT reconstruction.

To assess whether the preexisting medical conditions would influence the absolute EIT conductivity, we included the two subjects with electrolytes abnormities (n=18). We observed that the correlation value decreased from R = −0.69 (*p* = 0,003, n=16) to R= −0.21 (*p* = 0.4, n=18 (**Fig S2A**). We further excluded 2 subjects with anemia, and we noted that the correlation improved from R = −0.69 (*p* = 0,003, n=16) to R = −0.70 (*p* = 0.0049, n=14) (**Fig S2B**). These results were consistent with the impedimetric property underlying the composition of the liver in the setting of pre-existing conditions-associated electrolyte disturbance. In this case, the presence of leukemia, renal failure, and anemia disrupted the homeostasis of the organ systems; thus, altering the liver conductivity. Thus, our fundamental and experimental analyses pave the way for defining our exclusion criteria for future subject enrolment.

In summary, we enrolled overweight/obese subjects to undergo MRI scans and liver EIT measurements to generate the EIT conductivity map. We demonstrated that the increase in liver EIT conductivity is correlated with a decrease in MRI PDFF. As a corollary, we demonstrated that the 3-D EIT conductivity map also revealed the heterogeneous distribution of fatty gradient as evidenced by the 3-D MRI PDFF. Our correlation analyses supported that subject-specific EIT offers a non-invasive and portable method for the detection of hepatic fat infiltrate; thereby, proving a translational basis for developing liver EIT suitable for operator-independent, low-cost identification and monitoring of fatty liver disease.

## Methods

### Study Design

The recruitment of human subjects was conducted at the UCLA Center for Human Nutrition and Obesity in compliance with the UCLA Human Subjects Protection Committee. The study protocol (#15-001756) was approved by the UCLA Internal Review Board. All subjects provided written informed consent before participating in research procedures. We enrolled a total of 19 volunteers including, 15 females and 4 males, from 27 to 74 years old with a waist circumference from 91 cm to 141.5 cm and body mass index (BMI) from 25.5 to 46.8 kg/m^2^ (**Fig. 1**). Inclusion criteria for all subjects included an age range between 20-75 years, ability to travel for phlebotomy for whole blood collection, no prescription or over-the-counter medications for weight loss, and absence of alcohol consumption, and no weight change > 5 pounds in the previous 3 months. All subjects must be able to follow instructions and to consent. Exclusion criteria for all subjects included coronary artery disease on medications, claustrophobia, previous liver cancer, liver surgery, alcoholism (DSM-5 criteria: alcohol abuse or dependence), metallic implants or other factors hazardous to the MRI scanner as per the MRI safety guidelines, and body weight > 300 pounds (weight and size restrictions for undergoing MRI). Note that an MRI scan was performed to establish PDFF for quantifying obesity-associated fatty liver. Clinical demographic and physical characteristics of human subjects were collected in terms of gender, BMI (kg·m^−2^), age (years), waist circumference (cm), height (cm), and weight (kg) (**Table 2**). Following enrollment and consent, the subjects underwent a 30-min liver MRI scan, including multi-echo imaging for mapping the proton density fat fraction (PDFF) (**Fig. 6**). Next, EIT measurement was acquired by placing 32 electrodes to the upper abdominal region as indicated by the fiduciary markers immediately following the MRI scan (**Fig. 6A**). A pair of electrodes was used to inject the AC current to the abdomen, and the electrode array was used to record voltage by the pairwise algorithm **(Fig. 6B)**. Liver MRI provided the *a priori* knowledge of the boundary conditions needed for the EIT conductivity map reconstruction and PDFF **(Fig 6C-D)**. EIT conductivity map was reconstructed to distinguish the liver conductivity gradient from other tissues or organs **(Fig. 6E)**. Finally, subject-specific EIT (conductivity map) was compared with the corresponding MRI PDFF **(Fig. 6E)**.

**Figure 6.**
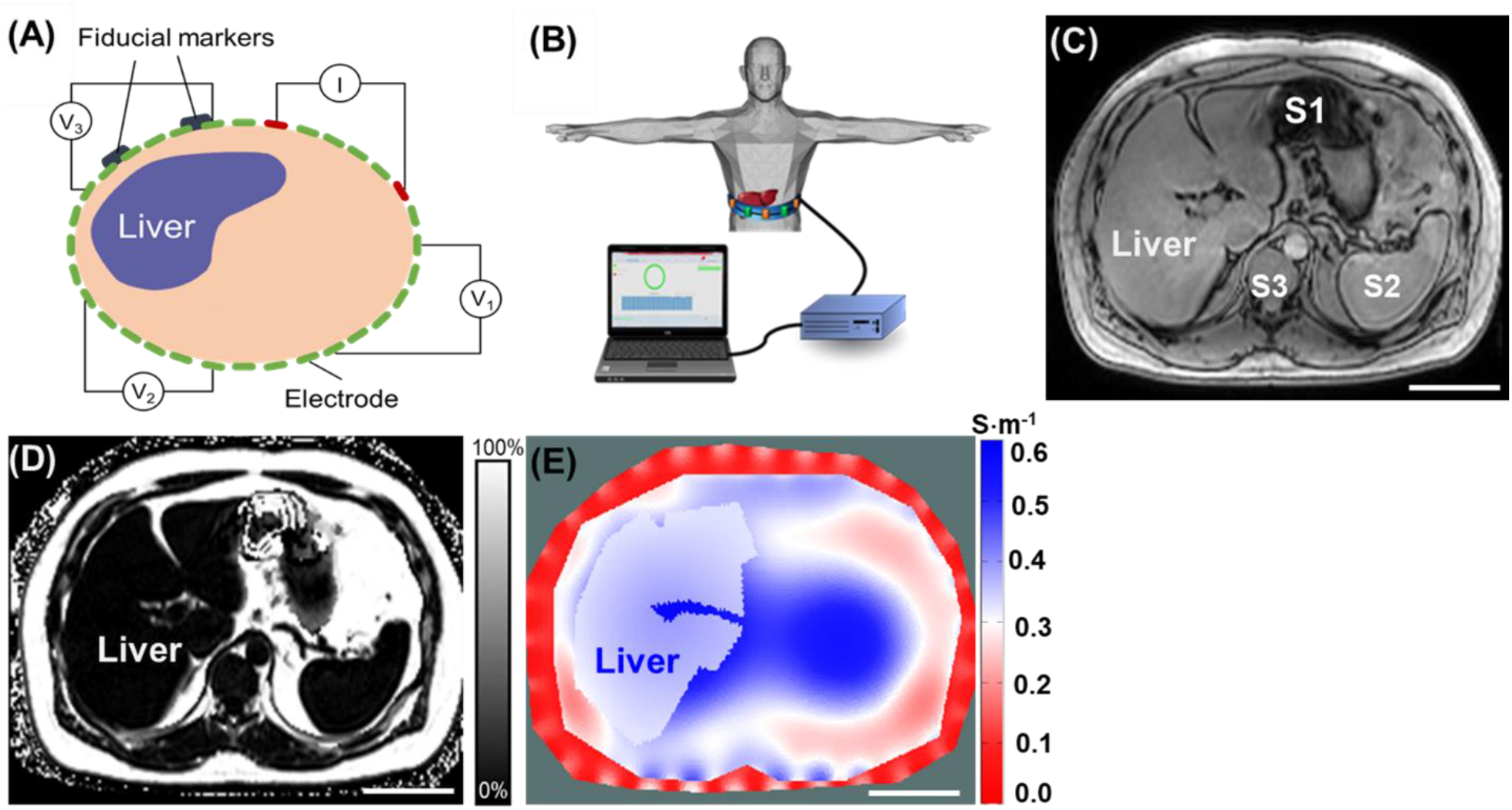
Schematic of EIT measurement, reconstruction, and 2-D representation. **(A)** Schematic illustrates circumferential electrode placement around the abdomen for pairwise voltage measurements. The fiducial markers indicate the anatomic level at which the multi-electrode array was circumferentially positioned for liver EIT measurements. **(B)** Thirty-two electrodes were adhered to the abdomen, as indicated by the fiducial markers. The recorded voltage signals were input to a signal adaptor and the data acquisition channels for EIT measurements. A representative MRI multi-echo image demarcates the boundary conditions **(C)** and PDFF map **(D)** for the abdomen, liver, stomach and spleen. S1: Stomach, S2: Spleen, S3: Spine. **(E)** A representative 2-D EIT image reveals the conductivity distribution. Scale bar: 8 cm.

### Determination of Liver MRI proton-density fat fraction PDFF

Non-contrast-enhanced abdominal MRI scans were performed on a 3-Tesla system (Skyra or Prisma, Siemens, Erlangen, Germany) using a body array and a spine array coils. The protocol included breath-held anatomical scouts, a breath-held T2-weighted 2-D multi-slice half-Fourier single-shot turbo spin-echo (HASTE) sequence, and a breath-held 3-D multi-echo gradient-echo sequence (TE = 1.23, 2.46, 3.69, 4.92, 6.15, 7.38 ms; TR = 8.94 ms, flip angle = 4 deg, typical field of view = 400 × 350 × 256 mm^3^, typical matrix size = 192 × 168 × 64, parallel imaging factor = 4, typical scan time = 19 sec) to quantify PDFF. Scanner software (LiverLab, Siemens, Erlangen, Germany), which utilized a multi-peak fat spectral model with single R_2_* for multi-step signal fitting, was used to calculate PDFF^50^. The MRI images and PDFF maps were saved in DICOM format and downloaded from the scanner for analyses.

To ensure alignment of the subsequent EIT slice position to a corresponding mid-liver MRI slice, we affixed two to three MRI-visible fiducial markers (MR-SPOT 122, Beekley Medical, Bristol, CT) to the skin above the expected mid-liver region prior to performing the MRI scan (**Fig 6A**). The positioning of the fiducial markers was examined on the anatomical scouts. If needed, the MRI technologist would re-position the fiducial markers on the subject’s abdomen and re-acquire the scouts. At least one adjustment would be required, and this entire alignment required less than 3 minutes.

The echo 1 (TE=1.23) magnitude images from the 3-D multi-echo gradient-echo sequence were used for contouring the body and the liver to create a 3-D anatomy model. An axial slice in the MRI PDFF maps that contained MRI-visible fiducial markers was selected for analysis. Five circular regions of interest (ROIs) with an area of 5 mm^2^ were delineated in the slice with fiducial markers by a trained researcher to avoid blood vessels, bile ducts, and imaging artifacts, and at least 1-2 cm away from the liver capsule. The mean PDFF from the ROIs (0-100%) was reported for each subject.

### Theoretical Framework for EIT reconstruction (EIDORS)

The EIT imaging reconstruction was implemented as previously described^11^. Following the injection of a known current to the abdomen, an EIT conductivity map across the abdomen was reconstructed with a set of voltages recorded by with an electrode array placed on the surface of the upper abdomen (see **Fig S3**)^7^. With a *priori* knowledge of the target object (liver), the geometric boundary conditions were established with a high degree of precision to mitigate instability inherent from the ill-posed EIT inverse problem^51^(**Fig. 6D**), and the solution was obtained by using a regularized Gauss-Newton (GN) type solver(**Fig. S3**).

The Gauss-Newton (GN) type solver calculates the conductivity by minimizing ∅, the l2 norm (the square root of the sum of the squares of the values) of the difference between the measured voltage *V_o_*, and a function of the conductivity *f*(σ):

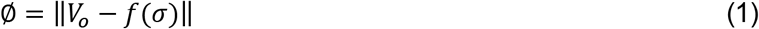

where *f*(σ) is considered to be the “forward problem” derived from the Laplace equations:

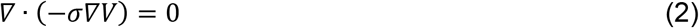

By taking the first order Taylor series expansion of ∅:

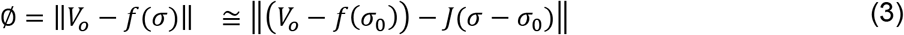

where *σ*_0_ is a reference conductivity value, and *J* is the Jacobian matrix of our inverse problem.

By setting 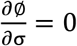, we minimized ∅ and obtained *σ* as follows:

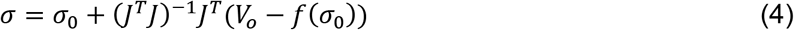

Equation (4) is an unconstrained GN form of the inverse problem. Due to the ill-posed nature of the EIT inverse problem, achieving a converged solution from this unconstrained GN form is challenging. The solution *σ* is highly sensitive to perturbations in voltage (*V*) measurement which means a small noise in *V* leads to instability in the final solution. A general method to mitigate the issue is to introduce a constraint term that sways the solution towards the preferred solution:

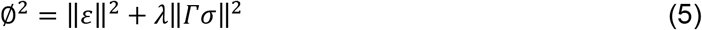

To balance the tradeoff between fitting the error and constraining the solution from the undesired properties, we incorporated a constraint term, *λ*||Гσ||^2^, to the objective function and the resulted form is commonly known as the Tikhonov Regularization. The coefficient, *λ*, is the regularization parameter that suppresses the conductivity spikes in the solution space.

With a *priori* conductivity within a similar area, the term, *Г*, was introduced as a “weighted” Laplacian operator that enables us to adjust more properties of the conductivity and suppress the non-smooth regions. Akin to the present work, this strategy is useful in medical imaging, where a *priori* anatomic information of individual organs was obtained from MRI multi-echo sequence and integrated with the EIT solutions. By applying the regulation term to equation (2), we generated the solution as follows:

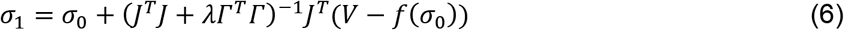

To obtain the absolute conductivity mapping, we adopted an iterated approach by first assuming an arbitrary conductivity, *σ*_0_, which is used to calculate *J*, *Г*, and *f*(*σ*_0_). From equation (6), we calculated a new conductivity value set, *σ*_1_ to generate a new set of *J*, *Г* and *f*(*σ*_1_). The iteration continued until the difference between *σ*_*n*_ and *σ*_*n*−1_ reached a minimally desired value.

In this study, we adopted the online open-source software suite EIDORS (version 3.8) for EIT image reconstruction. An inverse finite element model was aided with a mesh generator (Netgen) for the reconstruction of liver EIT image. Rather than using a presumed geometry for the finite element model, we combined the MRI multi-echo images-acquired geometric information with the multi-electrode-measured voltage data to reconstruct the liver EIT conductivity map. As a result, the computational errors from the variations in the geometry of the abdomen of different subjects were reduced.

### EIT measurement and reconstruction for fat infiltrate

Following MRI scans, the subjects underwent EIT measurement. The MRI-visible fiducial markers on the abdomen facilitated the circumferential positioning of EIT electrodes. Next, the subjects were instructed to perform the same breath-holds (e.g., end inspiration) as they did for the MRI scan to ensure that the EIT slice matched with the level of the mid-liver MRI slice. Electrical measurement and data acquisition were conducted using the Swisstom EIT Pioneer Set (Swisstom AG, Switzerland). An array of disposable surface electrocardiogram electrodes (Covidien, Ireland) was attached to the skin of the subject, and each of them was connected to one of the 32 data acquisition channels of the Swisstom system, which was interfaced with the controlling computer via a separate module (**Fig. 6A-B**). The AC currents with programmable magnitude from 2-4 mA were injected to the upper abdomen at 50 kHz and 250 kHz respectively through the selected channels, and the resulting voltage responses were recorded by a separate pair of electrodes. A “skipping 4” pattern was used for current injection and voltage recording^52^ (**Fig. 6A**). Based on the anatomy of the liver, we established both 2-D and 3-D forward models for EIT reconstruction using the EIDORS library. The acquired voltage data were used to calculate 2-D and 3-D conductivity distribution. Following EIT reconstruction of the liver conductivity maps, we compared the liver conductivity (S·m^−1^) and MRI PDFF with the subject-specific demographics, and we generated the confidence interval plots to demonstrate the correlation between liver EIT conductivity and MRI PDFF.

### Statistical Analysis

We performed correlation analyses in the context of the subjects’ demographics (n=19) (such as chronic lymphocytic leukemia and renal failure). We excluded one liver EIT measurement due to the malfunction of electrodes. We compared the differences in correlation values between the presence (n= 18) and the absence of pre-existing medical conditions that could disturb the circulation and tissue electrolytes (n=16). The correlation between liver conductivity and MRI PDFF was assessed by Pearson’s correlation analysis and the Bonferroni correction for multi-testing. The comparison between EIT and demographics, and between MRI PDFF and demographics, were analyzed for statistically significant coefficients and 95% confidence interval using Pearson’s correlation analysis in *R*.

## Supplementary Materials

**Fig. S1**. 3-D MRI PDFF mapping vs. 3-D EIT image.

**Fig. S2**. Sub-analysis of EIT liver conductivity vs. MRI PDFF for all subjects and additional exclusion of anemic subjects.

**Fig. S3**. Schematic flow of EIT reconstruction.

**Table S1**. Conductivities of human tissue.

---

**Supplementary Figure 1.**
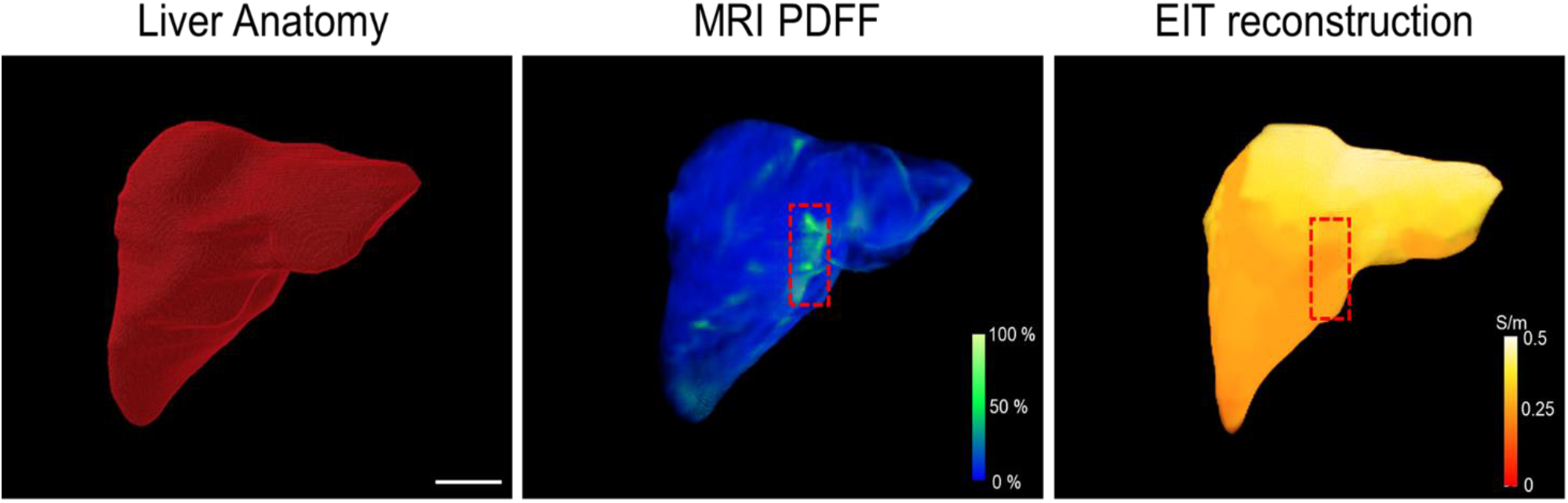
3-D MRI PDFF mapping vs. 3-D EIT image. **(A)** The representative 3-D liver boundary condition was established following segmentation of the MRI multi-echo imaging. **(B)** 3-D MRI PDFF mapping reveals a heterogeneous distribution of MRI PDFF. The red dashed box highlights the region with a relatively high fat fraction. **(C)** 3-D EIT image unveils the heterogeneous gradient of conductivity. The dash red box is consistent with that of MRI PDFF mapping. Thus, the 3-D comparison between MRI multi-echo imaging and EIT image further supports the correlation between MRI fat fraction and EIT conductivity. Scale bar: 5 cm.

**Supplementary Figure 2.**
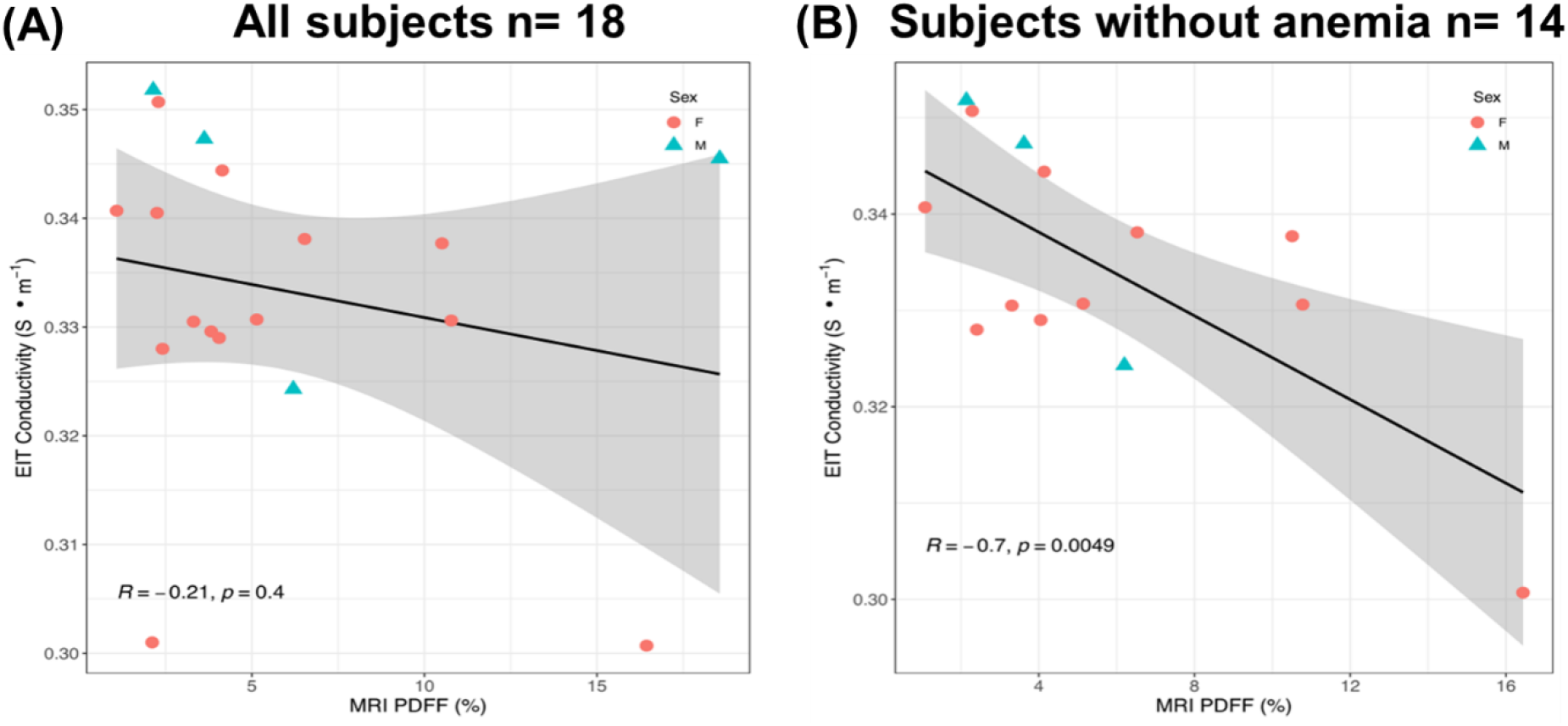
Sub-analysis of EIT liver conductivity vs. MRI PDFF for all subjects and additional exclusion of anemic subjects. **(A)** The negative correlation between EIT conductivity and MRI PDFF was reduced to R=-0.21 in the presence of preexisting medical conditions implicated in disturbing tissue electrolytes (*p* = 0.4, n= 18). **(B)** The correlation between EIT liver vs. MRI PDFF was increased to R = −0.70 in the absence of anemia subjects (*p* = 0.0049, n=14). The shaded areas reflect the 95% confidence intervals of the linear slopes.

**Supplementary Figure 3.**
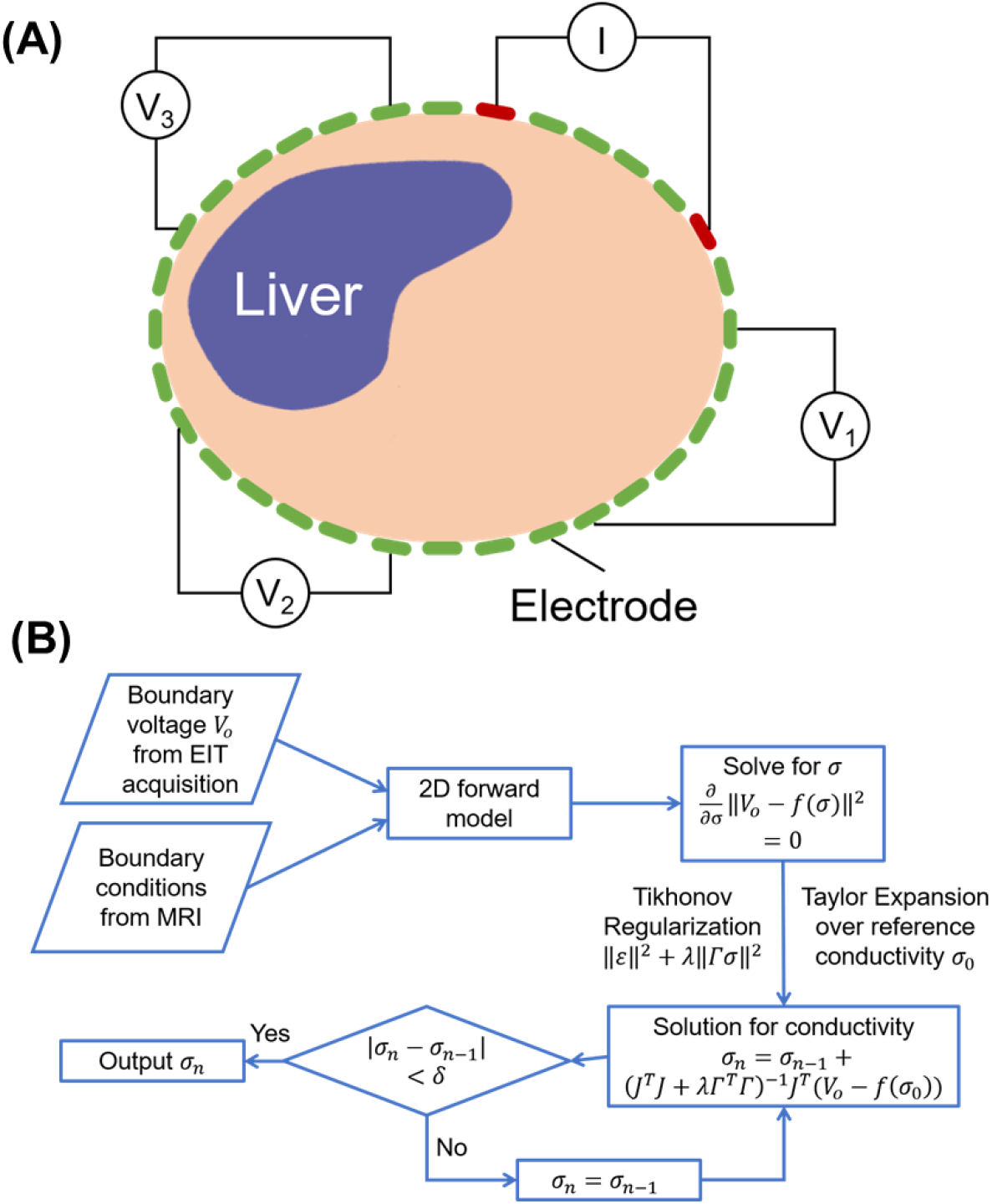
Schematic flow of EIT reconstruction. **(A)**“Skipping 4” pattern was used for both current injection and voltage acquisition. There were 4 electrodes separating each pair of stimulating and detecting electrodes. **(B)** EIT reconstruction was established by solving the inverse problem via a regularized Gauss-Newton (GN) type solver.

**Supplementary Table 1.** Conductivities of human tissues at 50 kHz [S·m^−1^].

| Tissue | $\text{S}\cdot\text{m}^{-1}$ | Tissue | $\text{S}\cdot\text{m}^{-1}$ |
| --- | --- | --- | --- |
| liver | 0.07 | fat | 0.04 |
| lung | 0.14 | muscle | 0.35 |
| heart | 0.10 | bone marrow | 0.06 |
| kidney | 0.10 | skin | 0.10 |
| intestine | 0.35 | blood | 0.70 |
| stomach | 0.50 | cartilage | 0.18 |

## Notes

### Competing Interest Statement

The authors have declared no competing interest.

